# A taxogenomics approach uncovers a new genus in the phylum Placozoa

**DOI:** 10.1101/202119

**Authors:** Michael Eitel, Warren R. Francis, Hans-Jürgen Osigus, Stefan Krebs, Sergio Vargas, Helmut Blum, Gray A. Williams, Bernd Schierwater, Gert Wörheide

## Abstract

The Placozoa [1] is a monotypic phylum of non-bilaterian marine animals. Its only species, *Trichoplax adhaerens*, was described in 1883 [2], Despite the worldwide distribution of placozoans [3–6], morphological differences are lacking among isolates from different geographic areas and, consequently, no other species in this phylum has been described and accepted for more than 130 years. However, recent single-gene studies on the genetic diversity of this “species” have revealed deeply divergent lineages of, as yet, undefined taxonomic ranks [3,5,6], Since single genes are not considered sufficient to define species [7], a whole nuclear genome comparison appears the most appropriate approach to determine relationships between placozoan lineages. Such a “taxogenomics” approach can help discover and diagnose potential additional species and, therefore, develop a much-needed, more robust, taxonomic framework for this phylum. To achieve this we sequenced the genome of a placozoan lineage isolated from Hong Kong (lineage H13), which is distantly related to *T. adhaerens* [6]. The 87 megabase genome assembly contains 12,010 genes. Comparison to the *T. adhaerens* genome [8] identified an average protein distance of 24.4% in more than 2,700 screened one-to-one orthologs, similar to levels observed between the chordate classes mammals and birds. Genome rearrangements are commonplace and >25% of genes are not collinear (i.e. they are not in the same order in the two genomes). Finally, a multi-gene distance comparison with other non-bilaterian phyla indicate genus level differences to *T. adhaerens*. These data highlight the large genomic diversity within the Placozoa and justifies the designation of lineage HI3 as a new species, *Xxxxxxxxx yyyyyyyyyyyyy*^*1*^ gen. et spaec. nov., now the second described placozoan species and the first in a new genus. Phylogenomic analyses furthermore supports a robust placement of the Placozoa as sister to a cnidarian-bilaterian clade.

## Results and Discussion

### Adding a new placozoan genome and improving the *T. adhaerens* genome annotation

Based on mitochondrial 16S rDNA analyses, the placozoan lineage HI3 is among the most distantly related lineage to *T. adhaerens* [6], whose nuclear genome has been sequenced previously [8]. We hypothesized that the substantial 16S rDNA divergence could also be reflected on the whole-genome scale and, therefore, targeted lineage HI3 for nuclear genome sequencing. To assemble the genome of lineage H13 – a new species described here called *Xxxxxxxxx yyyyyyyyyyyyy* gen. et spec. nov. (see species description in Methods) – we generated 24 Gb of paired-end reads and 320 Mb of Moleculo (Illumina Artificial Long Synthetic) reads. Our final, highly complete, 87 megabase assembly contained 669 high-quality and contamination filtered contigs with an gap-free N50 of 407 kb (Table S1; Figures S1-S3), seven megabases smaller than the *T. adhaerens* contig assembly. The overall calculated genome heterozygosity was 1.6% (based on SNP counts, see Table S2).

We annotated the genome with a combination of 15.3 Gb of RNAseq and *ab initio* methods to yield 12,010 genes (Table S1 and Supplemental Information). A high percentage of raw reads mapped back to the genome (Table S3) and between 89-93% of the 978 genes in the Metazoa BUSCO v2.0 dataset were identified in the different annotation sets (Table S4). Together this suggests an almost complete assembly and annotation, where more than 96% of the annotated genes in the *X. yyyyyyyyyyyyy* genome were expressed in adult animals. In our gene set, *X. yyyyyyyyyyyyy* had 490 more genes than the 11,520 genes reported in the original *T. adhaerens* annotation. We re-annotated *T. adhaerens* with AUGUSTUS and found an additional 1,001 proteins and also managed to complete formerly partial proteins. This approach added 4.4 Mb of exons to the *T. adhaerens* annotation, an increase of 28% of exonic base pairs to the original annotation. The new *T. adhaerens* annotation now has 511 more genes than *X. yyyyyyyyyyyyy*, which accounts for some portion of the size difference between the two genomes.

### Large-scale genomic distance analysis identifies large genetic divergence between *X. yyyyyyyyyyyyy* and *T. adhaerens*

The roughly 4x coverage of the genome with long Moleculo reads (N50 of 5.4kb) allowed the assembly of large haplocontigs (i.e. contigs representing both haplotypes of the genome). This phasing information for large parts of the genome facilitated the isolation of both full-length alleles at 2,720 loci after a highly stringent filtering procedure. Our thorough filtering allowed the confident grouping of orthologous alleles, except in rare cases, when two fundamental conditions were met, namely: (i) recently duplicated genes with highly similar sequences (even in introns) that fall below the filtering cutoff, and (ii) the true orthologous allele was missing in the genome assembly. Since the assembly is almost complete, the proportion of rare false positive alleles should be negligible. In addition, we identified and carefully validated (see Methods) orthologous sequences between *X. yyyyyyyyyyyyy* and *T. adhaerens* for these 2,720 loci. We are, therefore, confident that we used only true alleles as well as interspecific orthologs for the 2,720 loci in our sequence divergence analyses.

Between the two*X. yyyyyyyyyyyyy* alleles genetic distance ranged from 0.0 to 13.2% (mean 1.1% ±1.0) for proteins, and 0.0 to 9.5% (mean 1.0% ±0.5) for coding sequences, respectively, whilst between *X. yyyyyyyyyyyyy* and *T. adhaerens* genetic distance ranged from 0. 0 to 72.4% (mean 24.4% ±11.3) for proteins, and 5.1 to 55.5 (mean 24.5% ± 6.4) for coding sequences, respectively (Figure 1C). Most genes showing a high variation at the allelic level in *X. yyyyyyyyyyyyy* were also highly different between the species. To assess if certain genes are under positive (diversifying) selection, indicative of functional evolution, we calculated the ratio of nonsynonymous to synonymous nucleotide substitutions (dN/dS ratio, e.g. [9]) for each *X. yyyyyyyyyyyyy* and *T. adhaerens* ortholog pair. Our results show that most orthologs (97%) are under strong purifying selection (dN/dS <0.5). One might hypothesize that a strong purifying selection pressure is the reason for the phenotypic stasis we see in modern placozoans. More placozoan genomes across the diversity in the phylum are clearly needed to further test this hypothesis. Despite this strong tendency towards purifying selection, a high proportion of orthologs (49%) showed larger protein distance than coding sequence distance and therefore an accumulation of double or triple mutations per codon that led to amino acid substitutions. The reason for this pattern is unclear, and needs further investigation.

Only three of the 2,720 orthologs (0.1%) have dN/dS ratios slightly > 1, indicating positive selection (Supplemental Information; see Figure S4 for an estimate of mutation saturation in codons). The best BLAST hits of those three positively selected genes to Human Uniprots were SRF, SRN2, and IKKA, which are involved in transcription regulation, mRNA splicing and NF-kappa-B signaling, respectively. The function of these proteins in placozoans, however, still has to be studied in detail.

**Figure 1.**
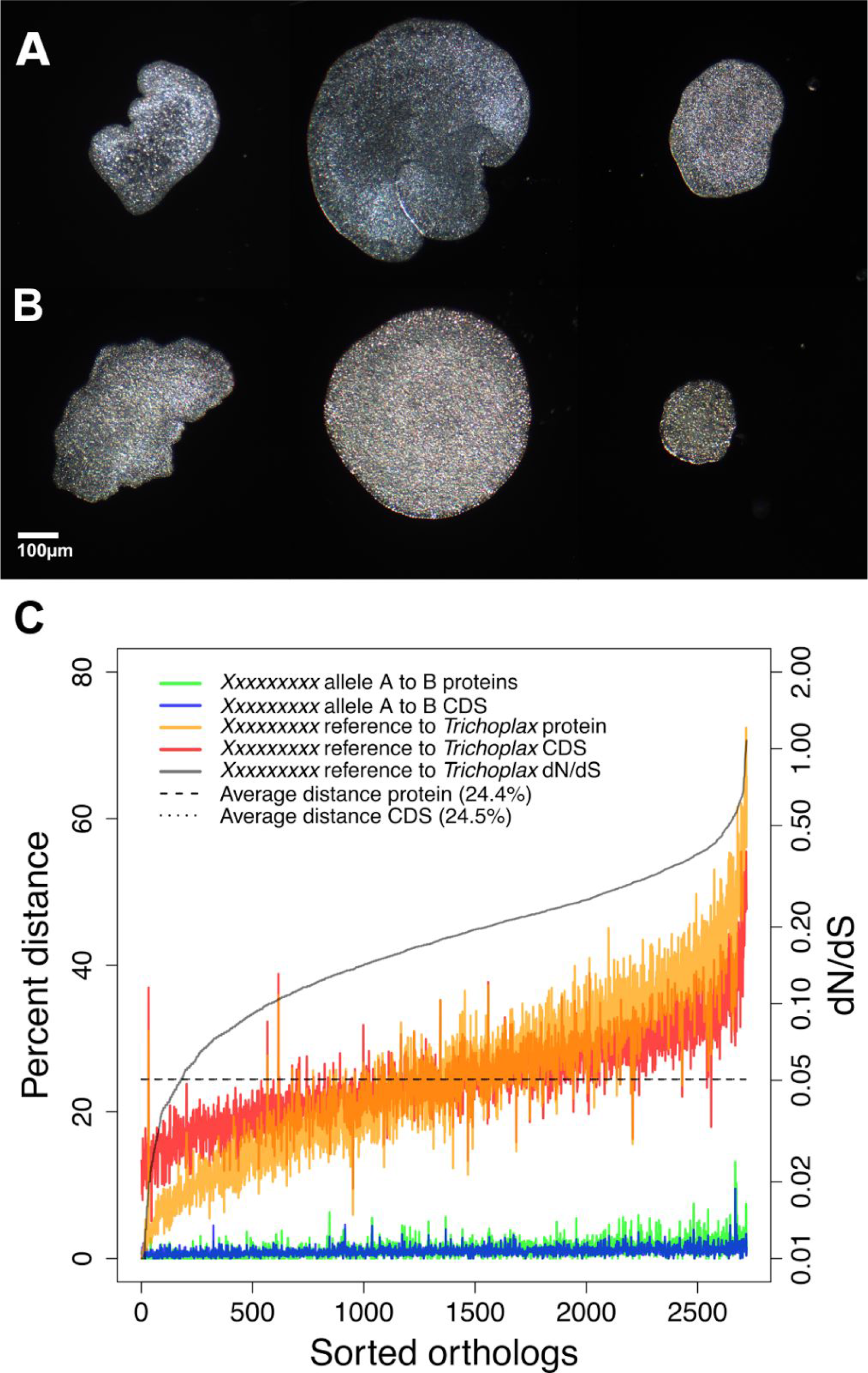
Live placozoan specimens and genetic distances. Light microscopy images of three (left to right) live *Trichoplax adhaerens* (**A**) and *Xxxxxxxxx yyyyyyyyyyyyy* (**B**) specimens. General body plans are identical for placozoans and at the same time intraspecific shape plasticity can be high, which prevents definition of reliable morphological characters. Scale bar applies to all images. (**C**) Pairwise allelic (blue, green line) and interspecific (red, orange line) distances for 2,720 orthologous genes. A large fraction of orthologs have larger protein than coding sequence (CDS) distance, but only three of these are in fact positively selected (reflected by dN/dS ratios > 1, gray line). Orthologs are sorted by increasing dN/dS.

### Genomic rearrangements are commonplace between the *X. yyyyyyyyyyyyy* and *T. adhaerens* genomes

Moleculo reads also enabled us to assemble very large reference contigs, the largest being over 2 Mb. We compared the organization of genes in *X. yyyyyyyyyyyyy* to the ten largest scaffolds in the *T. adhaerens* genome (size range 2.4-13.2 Mb; accounting for 66% of the *T. adhaerens* assembly). We found 144 contigs > 100 kb from *X. yyyyyyyyyyyyy* that aligned to these ten scaffolds, accounting for 69% of the *X. yyyyyyyyyyyyy* assembly (Figure 2). Mean gene collinearity (i.e. the same genes in the same direction) in this reduced genome representation was in the range of 69.5% to 78.8% (mean 73.6%±5.5; see Table S5). The mean number of genes per block was 33.8 (±25.2) in the reduced set and 33.9 (±24.7) when comparing full genomes, which indicates that the reduced set is representative for both full genomes (see Figure S5).

**Figure 2.**
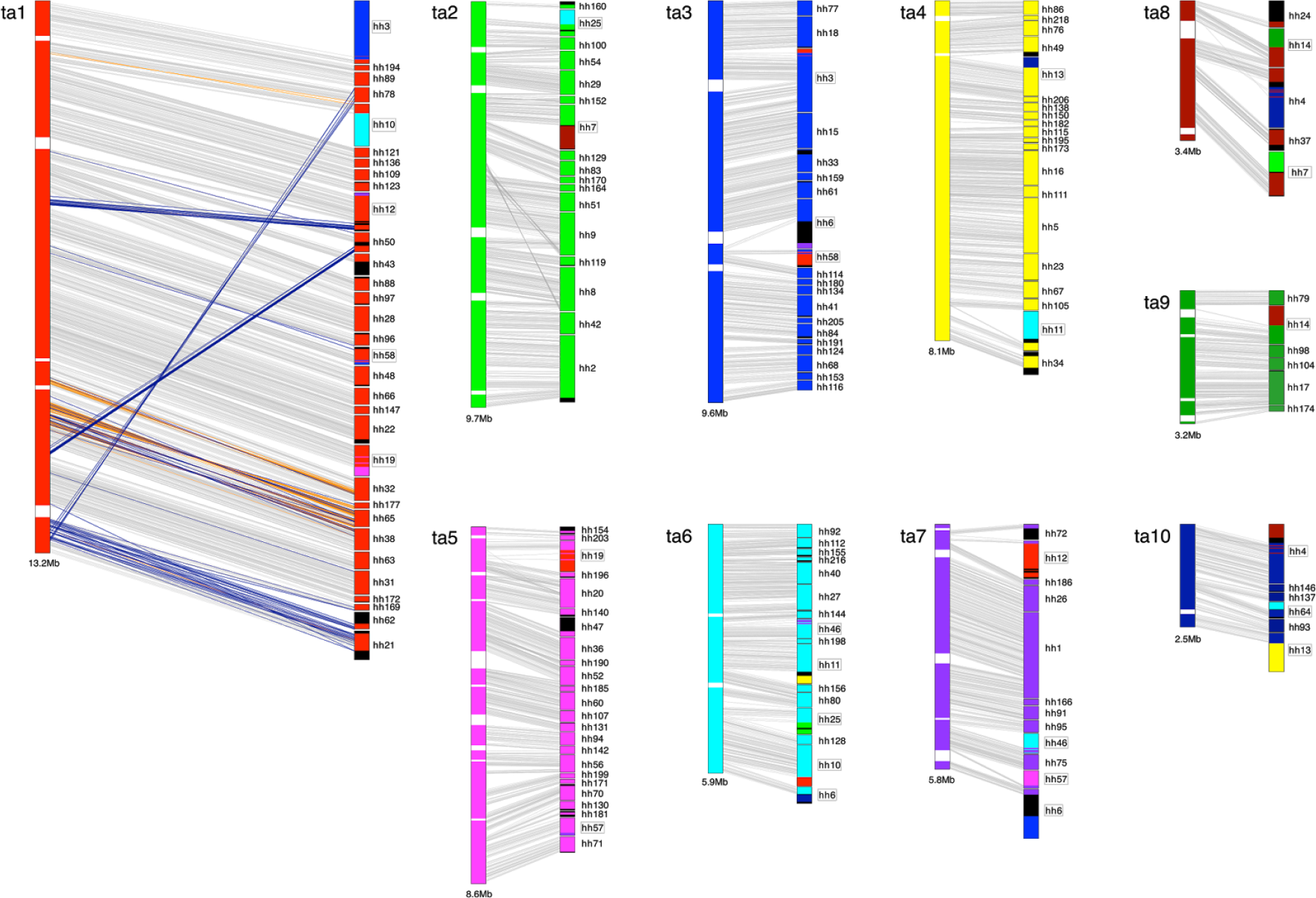
Synteny between *Xxxxxxxxx yyyyyyyyyyyyy* and Trichoplax adhaerens. Scaled schematic drawings of the ten longest *T. adhaerens* scaffolds on the left (tal-talO) and matching *X. yyyyyyyyyyyyy* contigs (hh) on the right. Each line represents one gene. While a general macro-synteny between the two placozoan species is present, 25% of the genes are translocated or inverted relative to the order of the respective *T. adhaerens* scaffold (orange and blue lines, respectively; illustrated for tal only). Often, entire blocks are translocated (different colors in boxed *X. yyyyyyyyyyyyy* contigs). Black stretches mark genomic regions not matching any of the ten *T. adhaerens* scaffolds while white stretches mark gaps in the *T. adhaerens* scaffolds.

Although much of the gene order is conserved between the two species, we counted 2,101 genes (out of the 8,260 genes in the ten scaffolds) that were inverted or translocated within the same scaffold relative to the order in the *T. adhaerens* scaffolds. These numbers seem low when compared to the fast evolving bilaterian genus *Drosophila* [10,11] or the even more extreme *Caenorhabditis* [12], but they are in the range of rearrangements found between mouse and human [13]. Comparison to Bilateria, however, might be misleading since rates of evolution are not directly comparable and genome rearrangement events might be more favoured in some bilaterian taxa due to intrinsic genomic traits such as transposon-induced rearrangement hotspots (e.g. [14]). Nonetheless, the high percentage of rearrangements between *T. adhaerens* and *X. yyyyyyyyyyyyy* adds further support to our taxonomic decisions, but adding more placozoan genomes to the analyses is essential for a more complete evaluation of how genome rearrangements can be used in placozoan taxonomy and systematics.

### A taxogenomic approach allows the description of a new placozoan species and genus

All internal Linnean ranks within the Placozoa are as yet undefined (e.g. [6]). Reliable diagnostic morphological characters, commonly used for defining species, are lacking in the Placozoa, despite efforts [15] to identify such, and reproductive isolation cannot, as yet, be tested [16–18], Thus, all present taxonomic definitions in this phylum must solely rely on diagnostic molecular characters. In other taxonomic groups (e.g. bacteria and archaea [19], protists [20,21], and fungi [22]), approaches and working models for purely sequence-based distinction of taxa have been proposed and are generally well established and widely accepted. In animals, such approaches are currently under development and have been used in rare cases to identify and describe cryptic species (e.g. [23]). Consequently, we here apply the genetic species concept *sensu* Baker & Bradley [24], which defines speciation as the accumulation of genetic changes in two lineages that depends on divergence in genes, the genome, and chromosome structure, to assess taxonomic relatedness and diagnose distinct placozoan species using a “taxogenomic” approach. We define taxogenomics as the integration of genomics into taxonomy (see also [25]). In addition to the placozoan genome structure variation (above), we compared the genomic variation across six different molecular sets of criteria between *X. yyyyyyyyyyyyy* and *T. adhaerens* to the variation in the three other non-bilaterian phyla: Cnidaria, Ctenophora, and Porifera (Figure 3A, Figures S6-S11). To achieve this, we used separate marker sets from different information sources (non-coding vs. protein coding genes) and origins (nuclear vs. mitochondrial genome) as criteria to evaluate congruence of these sets in a taxonomic framework. Marker sets included a nuclear protein set of 212 concatenated proteins (dataset 1: extended matrix from [26]; Tables S6-S8) as well as five selected genes with different substitution rates (nuclear 18S, 28S rDNA; mitochondrial 16S rDNA; mitochondrial proteins COl andNDl), all commonly used for DNA barcoding and molecular systematics.

Across individual markers, it appears that the phylogenetic ranks are most robust in the Cnidaria, where molecular variation corresponds well to classical taxonomy, in that higher ranks consistently correspond to greater distance between groups (Figures S6-S11). Measured distances for families within orders in Ctenophora and for genera within families in Porifera indicate that classical morphological taxonomies are incongruent with the calculated genetic distances (Figure 3A and Figures S6-S11). The internal phylogeny of these two phyla appears to be in urgent need of further re-evaluation with the inclusion of molecular data. The only non-bilaterian phylum with a consistent taxonomy, which is mirrored by genetic distances is, therefore, the Cnidaria. We consequently used genetic distances in the Cnidaria as an approximation and comparative guideline for the taxonomic classification of the new placozoan species.

**Figure 3.**
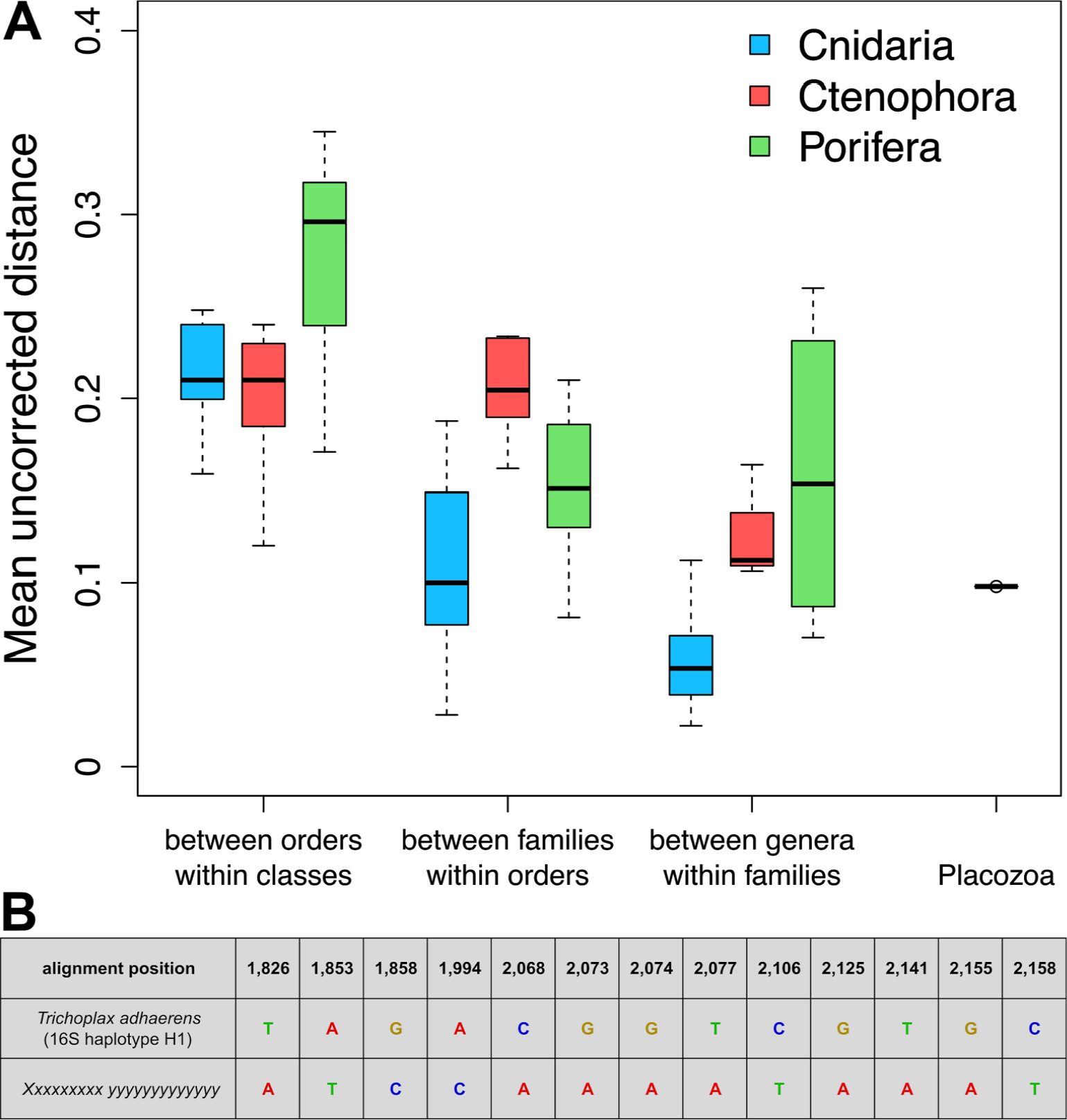
Calculated uncorrected pairwise genetic distances for 212 concatenated nuclear-encoded proteins (dataset 1). (**A**) Mean group distances for different taxonomic ranks in the non-bilaterian phyla Cnidaria, Ctenophora and Porifera. The interspecific protein distance of 11% between *Xxxxxxxxx yyyyyyyyyyyyy* and *Trichoplax adhaerens* (right) is comparable to mean group distances between genera within families in the Porifera and Ctenophora, respectively. With respect to the Cnidaria, the placozoan distance is even comparable to the mean group distance between families within orders. (**B**) Molecular diagnostics that characterise *Xxxxxxxxx yyyyyyyyyyyyy* based on differences in the alignment of the mitochondrial large ribosomal subunit (16S). See Figures S6-S10 for genetic distances in single marker genes and Figure S11 for a summary of all distances.

Genetic distances between *X. yyyyyyyyyyyyy* and *T. adhaerens* were higher than those for the Cnidaria in five of the six marker sets at the generic level, but lower at the family level in all cases (Figure S11, Table S9). Based on these results we cautiously place *X. yyyyyyyyyyyyy* as the first species of a new genus and assign *Xxxxxxxxx* gen. nov. to the Trichoplacidae fam. nov., a family including *Trichoplax* that we here define for the first time, since it was never formally assigned (albeit mentioned in [27]).

Among all markers, 16S rDNA appeared to be most variable among placozoans and other non-bilaterian phyla and the mean pairwise distance is closest to that calculated for the nuclear dataset in most cases (Figure S11). This marker also best mirrored classical taxonomy in the Porifera and Cnidaria (Figure S8; in Ctenophora 16S rDNA is highly derived and hard to identify [28]). According to these data, molecular diagnostics based on differences in the 16S rDNA appear to be suitable for current and future faster designation of species in the Placozoa, which is in agreement with previous results [5], Hence, we used sequence differences based on the 16S rDNA alignment as diagnostic characters in the species description of *Xxxxxxxxx yyyyyyyyyyyyy* to delimitate this new species from *Trichoplax adhaerens* (Figure 3B and species description in Methods).

### The *X. yyyyyyyyyyyyy* genome adds support to the phylogenetic placement of the Placozoa in the animal tree of life

Recent discussions about the phylogenetic position of placozoans have been based on the *T. adhaerens* genome. A better sampling of the placozoan genomic diversity is, however, needed [29] to address the current dispute over the phylogenetic relationships between early-branching metazoan phyla [30–32] and the placement of the Placozoa in the metazoan tree of life. In this context, it is important to first assess if adding another placozoan species would break up the long placozoan branch, because the inclusion of a single representative of a clade with a very long terminal branch, or fast-evolving taxa that can have random amino acid sequence similarities, may result in erroneous groupings in a phylogeny (so-called “long-branch attraction artefacts”) [32,33]. To address these questions, we generated a highly (taxa) condensed version of the full protein matrix (termed dataset 2 with less than 11% missing characters) and, additionally, created a Dayhoff 6-state recoded matrix [34] of this second set to reduce amino acid compositional heterogeneity, which is also known to be a source of phylogenetic error [35]. Phylogenetic analyses were performed on these two matrices (protein and Dayhoff-6 recoded), using the CAT-GTR model in PhyloBayes-MPI vl.7 [36]. The resulting trees suggest a sister group relationship of the Placozoa to a Cnidaria+Bilateria clade (Figure 4; Figures S12 and S13), in agreement with some previous findings [8,26,30,32,37,38] and with a recent study using a large geneset and intense quality controls [32].

**Figure 4.**
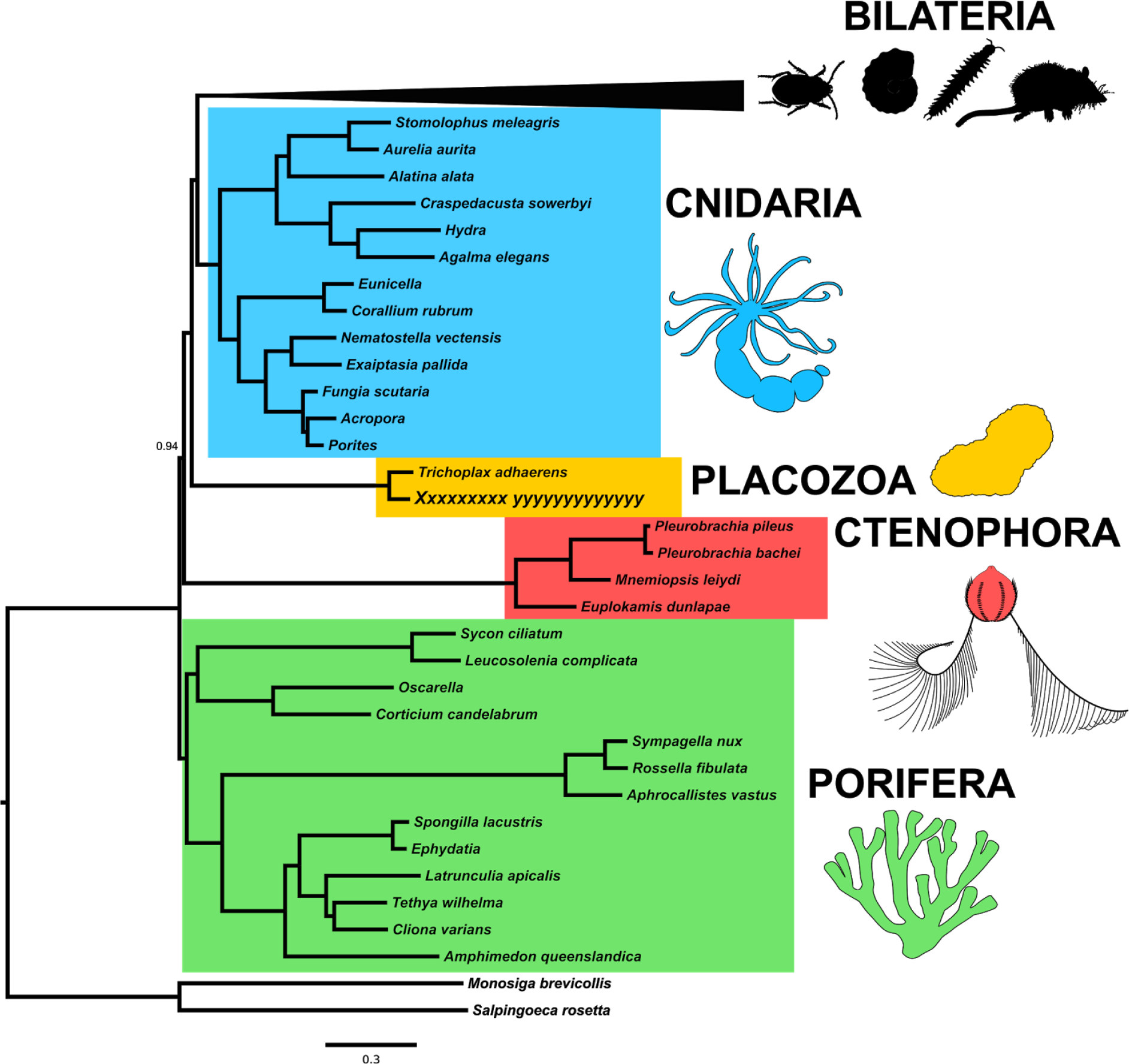
Phylogenetic tree based on the PhyloBayes analysis of dataset 2. According to this analysis, the Placozoa are sister to a cnidarian+bilaterian clade. Node supports are 1.0 unless otherwise noted. See Figure S12 and Figure S13 for the full trees using the protein and Dayhoff-recoded matrix.

### Conclusions

We have shown that a large and, as yet, insufficiently explored genomic diversity exists within the phylum Placozoa. Using a taxogenomic approach, based on molecular data only, we here discovered and described the only second species in the Placozoa. Future research efforts, including genome sequencing of additional placozoan lineages, will likely help to establish a broader and more robust systematic framework in the Placozoa and provide further insights into the mechanisms of speciation in this enigmatic marine phylum. The inclusion of a second placozoan species into phylogenomic analyses already does split the long placozoan branch to some extent, but also here more genomes from diverse placozoan species are needed to further manifest the phylogenetic placement of the Placozoa among the non-bilaterian animals and to improve our understanding of the evolution and diversity of placozoans.

## Author Contributions

M.E., W.R.F., B.S. and G.W. designed the study; M.E. and G.A.W. collected samples and cultured animals; M.E. prepared DNA for genome sequencing; H.J.O. and B.S. generated RNAseq data; S.K. and H.B. generated genomic libraries and performed sequencing; M.E. assembled the genome and transcriptomes, performed annotation, allele and ortholog assignments, and distance calculations; M.E. and W.R.F generated the functional annotation table; M.E. built and quality controlled the supermatrix for phylogenetic analyses with contribution from W.R.F; W.R.F. wrote Python scripts; S.V. and G.W. provided, and S.V. and W.R.F. analysed, sponge transcriptome and proteome data; G.W. and M.E. performed phylogenetic analyses. M.E., H.J.O. and W.R.F. generated the main figures; M.E. and W.R.F. wrote the first draft of the manuscript. All authors amended the manuscript and approved the final version.

## Acknowledgments

We thank Prof Kenneth Mei Yee Leung for identifying a meaningful genus name and Ms Cecily Law for assistance in animal culturing. We thank Dr Johanna Taylor Cannon for providing untrimmed alignments for the 212 nuclear proteins. Prof John Hooper is acknowledged for assistance in species description formalities. We thank Dr Nora Dotzler for preparing the *Xxxxxxxxx yyyyyyyyyyyyy* holotype specimen. This research benefitted from funding by LMU Munich’s Institutional Strategy LMUexcellent within the framework of the German Excellence Initiative to G.W. and H.B., two German Science Foundation (DFG) research grants (SCHI 277/26-1, SCHI 277/29-1) to B.S., and by a Small Project Fund of The University of Hong Kong provided to G.A.W. and M.E.․ H.J.O. acknowledges a doctoral fellowship of the Studienstiftung des deutschen Volkes. The German Academic Exchange service (DAAD) is acknowledged for a postdoctoral research fellowship to M.E. that supported specimens collection and establishment of cultures.

## METHODS

### Formal Taxonomic Diagnosis

*Phylum:* Placozoa, Grell 1971

*Family:* Trichoplacidae, fam. nov., Eitel, Schierwater & Wörheide *Trichoplax adhaerens*, the only described species in the phylum, was never formally assigned to a family, which we here define as Trichoplacidae, fam. nov.

*Genus: Xxxxxxxxx*, gen. nov., Eitel, Schierwater & Worheide First species of the genus, *Xxxxxxxxx yyyyyyyyyyyyy*, spec. nov.

*Type species: Xxxxxxxxx yyyyyyyyyyyyy*, spec, nov, Eitel, Schierwater & Wörheide

*Etymology:* WILL BE AVAILABLE UPON FORMAL JOURNAL PUBLICATION.

*Species: Xxxxxxxxx yyyyyyyyyyyyy*, spec, nov., Eitel, Schierwater & Wörheide

*Diagnosis:* Gross and fine morphology is identical for placozoans. Diagnostic characters are thus defined by nucleotide differences in the mitochondrial large ribosomal subunit (16S). Full-length 16S sequences of *Trichoplax adhaerens* and *Xxxxxxxxx yyyyyyyyyyyyy* were aligned with MAFFT using the GINSI option and otherwise default settings. Ambiguously aligned 5’ and 3’ sequence ends were removed. The final alignment length was 2,498 nucleotides (including gaps). The region for diagnostic nucleotides was restricted to a highly variable region of the alignment (as previously shown for a large range of placozoan lineages, [3,5,6]). Molecular diagnostics for *X. yyyyyyyyyyyyy* are given in Figure 3B.

*Type locality:* A single specimen of *X. yyyyyyyyyyyyy* was isolated in the Ho Chung River close to a small mangrove patch at Heung Chung village (22.352728N 114.251733E) on June 6th 2012.

*Type specimen:* One specimen from a clonal lineage of *Xxxxxxxxx yyyyyyyyyyyyy* has been mounted and deposited at the Bayerische Staatssammlung für Paläontologie und Geologie in Miinchen, Germany, under voucher number SNSB-BSPG.GW30216. In addition 20 clonal individuals have been stored in ethanol as paratypes under voucher number SNSB-BSPG.GW30217 in addition to a DNA extraction under voucher number SN SB-B SPG. GW3 0218.

*Etymology:* WILL BE AVAILABLE UPON FORMAL JOURNAL PUBLICATION

### Animal Material

Two strains were used for this project: The ‘M2RS3-2’ strain was used for the DNA sequencing (the ‘DNA strain’) and the ‘M153E-2’ strain (the ‘RNA strain’) for the transcriptome. Both strains descend from a single placozoan individual each, that were isolated from mangroves/mangrove associates at two different sites in Hong Kong (SAR, China). The DNA strain was isolated from a dead mussel shell collected in the Ho Chung River close to a small mangrove patch at Heung Chung village (22.352728N 114.251733E) on June 6th 2012. The habitat undergoes daily changes in salt concentration and the salinity at collection was 20psu. The RNA strain was isolated from collection traps (for details on slide sampling see [4]) connected to a mangrove-associates (*Hibiscus*) and highshore mangrove (*Excoecaria*) trees at Tai Tam Tuk (22.244708N 114.221978E) on March 30th 2012. Both clonal cultures were cultured in 14cm glass Petri dishes as described [18] with a pure *Pyrenomonas helgolandii* algae culture (strain ID 28.87, Culture Collection of Algae, Georg-August-Universität Göttingen). The two different strains were used for DNA and RNA sequencing, respectively, to identify polymorphisms in these strains living in the same habitat but at two hydro-geographically distinct sampling sites (northeast vs. southeast Hong Kong).

### Genome sequencing and assembly

#### Short read sequencing

DNA was isolated as described [39] from roughly 1,000 healthy growing and clonally dividing individuals. 150 ng of genomic DNA was used to prepare an Illumina-compatible paired-end library with a nominal insert size of 250bp. All steps were done using the reagents from the Accel DNA 1S library preparation kit (Swift Biosciences, Ann Arbor, USA) following the manufacturer’s protocol. A total of 120,429,967 125-bp pairs were sequenced on an Illumina HiSeq l500. An initial read quality check in FastQC (http://www.bioinformatics.babraham.ac.uk/projects/fastqc/) identified a low quality stretch of the first 8bp in each read, which was clipped with Trimmomatic v0.35 [40] [added options: HEADCROP:8], Clipped reads were subsequently filtered using the BioLite v0.4.0 filtering tool [41] [added options: -q 28 -t 33 -a -b]. All reads with an average Phred Quality Score below 28 and/or reads with vector contamination were removed entirely without trimming. Quality filtering reduced the dataset to 103,388,888 2x117bp high quality reads (total 24.2Gb equalling ~277x genome coverage).

#### Molecido long read sequencing

Moleculo reads were prepared using the TruSeq^®^ Synthetic Long-Read DNA Library Prep kit following the manufacturer’s protocol (Illumina, San Diego, USA). A total of 500 ng high molecular weight genomic DNA was used as input for the library preparation. Two lanes of the barcoded library were sequenced on an Illumina HiSeql500 run and assembled using Illumina’s cloud-based service (BaseSpace Sequence Hub). A total of 83,688 Moleculo reads >500bp were generated with a N50 of 5.4kb, a peak at 8kb, and a total size of 320Mb. Trimming of low quality and vector regions was performed with Geneious R8 [42] [added options: error probability limit 0.01; maximum low quality bases 80; maximum ambiguities 4] and resulted in 79,974 high quality Moleculo reads >500bp (totalling 313Mb). Moleculo reads assembly in Geneious R8 [added options: minimum overlap of 400bp; 100% identical overlaps; no gaps allowed] resulted in 49,793 assembled sequences (contigs and singlet) with a N50 of 7.5kb (total 258Mb equalling ~2.9x genome coverage).

#### dipSPAdes hybrid assembly

A mixed read type assembly was performed with the SPAdes 3.5.0 package [43,44]. Filtered paired-end reads were error corrected within the assembly pipeline which consists of (1) error correction, (2) SPAdes haplocontig assembly) and (3) dipSPAdes haplocontig merging. The assembled Long Artificial Reads were input as ‘-trusted contigs’ [other added options: --cov-cutoff 10 --careful -k 39,49,59,69,79,89,99,109. dipSPAdes merging resulted in a total of 777 contigs >500bp].

#### Contamination screening

DipSPAdes haplocontigs were screened for bacterial contaminations by TBLASTN searches (evalue le-10) using proteins from the Candidatus *Midichloria mitochondrii* (order Rickettsiales) genome, the bacterial species most closely related to the previously identified *T. adhaerens* endosymbiont [45], In a second TBLASTN search we used plasmid-encoded proteins from all Rickettsiales genomes at NCBI (May 2016) to identify putative plasmid-associated contigs. All candidate bacterial chromosome and plasmid contigs (n=19) were re-BLASTed (BLASTN & TBLASTX) against complete Rickettsiales genomes to confirm the bacterial origin and were subsequently removed from the *X. yyyyyyyyyyyyy* nuclear genome assembly. The mitochondrial chromosome was further identified by BLASTN searches (evalue 1e-20) using the Placozoa sp. ‘Shirahama’ mitochondrial genome [46] (Genbank accession NC_015309.1) and also removed from the nuclear genome contigs.

#### Supercontig generation

After contaminant removal supercontig generation was performed. In the first place 50bp were clipped off from both ends of all dipSPAdes consensus contigs as the coverage towards the ends of contigs drops and errors might accumulate. After clipping contigs <500bp were removed. Remaining contigs were assembled in Geneious R8. To identify correct overlaps *ab initio* gene models were generated for the contigs before assembly with AUGUSTUS 3.0.3 [47], AUGUSTUS was trained online using the WebAUGUSTUS service (http://bioinf.uni-greifswald.de/webaugustus) using the clipped genomic contigs and a reduced set of Trinity transcripts (see below). This set only included “c0_gi_il” components of all transcripts and consisted of 33,708 transcripts. After the training AUGUSTUS was run with the resulting species parameter output [added options: species=placo_hl3, strand=both, genemodel=atleastone, codingseq=on, protein=on, cds=on, sample=100, keep_viterbi=true, alternatives-from-sampling=true, minexonintronprob=0.2, minmeanexonintronprob=0.5, maxtracks=10, GFF3=on, exonnames=on].

Settings used in the Geneious supercontig assembly were: 5kb minimum overlap, 2% maximum mismatch per contig, 2% maximum gaps per contig, 2000bp maximum single gap size (to account for larger indels), and 40bp word length. Overlapping contigs were checked in Geneious for identical exons/intron structure of predicted AUGUSTUS gene models in the overlap. In case of <100% overlap sequence identity one or both contigs were trimmed manually to keep a 100% identical overlap. Consensus supercontigs were then called in Geneious.

Even after the dipSPAdes merging step and the Geneious assembly some overlapping haplocontigs were identified by BLASTN of all against all contigs. Merging of these haplocontigs was performed with a second round of Geneious supercontig assembly with less stringent settings: 5kb minimum overlap, 25% maximum mismatch per read, 15% maximum gaps per read, 2000bp maximum single gap size, and 24bp word length. Overlapping contigs were checked again for identical AUGUSTUS gene models. In the case of missing annotation on both sequences, BLASTN searches of both haplocontigs were performed against all supercontigs. Haplocontigs were merged if both sequences hit itself or the overlapping haplocontig only. Trimming of overlaps was carried out as mentioned above. Supercontig consensus calling was done in Geneious with default settings. Overlapping contigs with insertions in one contig of up to 2kb were merged based on Moleculo read support. For this Moleculo reads were mapped to the supercontigs in Geneious in ‘low stringency’ mode.

A third Geneious assembly was performed to remove internal allelic redundant contigs, i.e. haplocontigs with full overlap to a supercontig. Low stringency settings for this final Geneious assembly were: 0.5kb minimum overlap, 25% maximum mismatch per read, 15% maximum gaps per read, 2000bp maximum single gap size, and 24bp word length. Both the fully overlapping (internal redundant) haplocontig as well as the partially overlapping contig were BLASTed (BLASTN) against all supercontigs to confirm matches of the full-length overlap in only two highly confident (1e-100) BLAST hits. In addition internal allelic contigs were confirmed by identical AUGUSTUS models on both alleles. Confirmed internal allelic (redundant) contigs were then removed.

This procedure ended in a genomic assembly of 669 gap-free supercontigs with a N50 of 407.8kb and a total of 87,194,036bp. These contigs are hereafter termed “reference contigs”. Additional scaffolding was not performed as Moleculo reads bridged most complex regions and no additional reads were available for further scaffolding. For *Xxxxxxxxx yyyyyyyyyyyyy* assembly and annotation statistics and a comparison to the *Trichoplax adhaerens* see Table S1 [48] [added options: -s -norna -a -inv -lcambig -source -html -gff -e hmmer & -small for softmasking] using the "*T. adhaerens*" reference of the Dfam database [49] for hmmer-based searches.

### Transcriptome sequencing and assembly

#### Library preparation and sequencing

RNA was extracted from the RNA strain in two batches of 100 clonal individuals each using standard phenol/chloroform extractions. RNA was shipped to the New York Genome Center (New York, NY, USA) for RNA quality check, library preparation and sequencing. Strand-specific libraries were prepared with 500ng total RNA using the TruSeq stranded mRNA V2 kit (Illumina, San Diego, USA). The nominal library insert size was 300bp. A total of 61,313,870 strand specific 125-bp RNA pairs (13.1Gb) were sequenced on an Illumina HiSeq2500.

#### Transcriptome assembly

Prior to Trinity assembly reads were quality checked in FastQC and filtered with BioLite 0.4.0 [added options: -q 25 -t 33 -a -b] keeping all reads with an average Phred Quality Score >25. This reduced the number to 57,237,523 high quality read pairs. Reads were assembled with Trinity v2.0.6 [50,51] [added options: --seqType fq --SS_lib_type RF --normalize_reads --trimmomatic --max_memory 50G]. A total of 124,155 transcripts were assembled with an N50 of 2,550bp and an average length of 1,506bp.

### Genome annotation

#### Genome-based transcript generation

Filtered (see above for Trinity assembly) strand-specific paired RNA reads were mapped to the hardmasked reference contigs with Tophat2 v2.1.0 [52] [added options: --library-type fr-firststrand]. The Tophat2 output bam file was used to run StringTie v1.2.2 [53] with default settings on the hardmasked reference contigs. Finally StringTie transcripts and predicted protein and encoded protein sequences were created with TransDecoder v2.1 [51] and default settings.

#### Ab initio *gene prediction*

The softmasked reference contigs were run in the BRAKER1 v1.9 [54] pipeline with default settings using the Tophat2 bam file of mapped RNAseq reads as guidance. BRAKER1 predicted 12,010 genes and 12,575 transcripts (Table S1).

#### Identification of unexpressed de novo gene models

To calculate the amount of unexpressed *de novo* BRAKER1 predicted proteins we identified their overlap with StringTie and Trinity transcripts using BEDtools *intersect* [added options: -s -v -f IE-4 -r]. Gene model IDs, extracted from the resulting table, were used to extract expressed (models with overlapping/coincident RNAseq-based transcripts) and non-expressed gene models from the BRAKER1 annotation GFF file. Of the 12,010 BRAKER1 genes only 422 (3.5%) were not expressed.

#### Functional annotation

We performed local BLASTX searches [55] of StringTie transcripts against (1) *T. adhaerens* reference proteins from NCBI, (2) Uniprot proteins (http://www.uniprot.org/. [56]), and (3) *X. yyyyyyyyyyyyy* predicted BRAKER1 proteins [added options in all cases: -evalue 1e-10 -max_target_seqs 2 -outfmt 6]. For BLAST searches the standalone BLAST+ suite v2.6 [57] was used. To identify domains in the *X. yyyyyyyyyyyyy* proteome we performed an HMMscan on the StringTie transcripts using Hidden Markov Models of Pfam-A release v30.0 [58] with HMMER v3.1b2 [59,60]. The resulting table (Supplemental Information) was used to generate a GFF3 annotation file of the domains based on the StringTie transcripts with a custom Python script (*pfam2gff.py*). A combined BLAST and pfam annotation table was created using a custom Python script (*collectannotationinfo.py*). tRNAs were predicted with tRNAscan-SE on the reference contigs with default settings and stored in an annotation GFF3 format.

### Genome coverage

A “lavalamp” kmer/GC plot was generated (Figure S1) to yield a high resolution plot of read counts per %GC and 31bp kmer coverage using the Jellyfish kmer counter and a set of custom Python scripts (*kmersorter.py* & *fastqdumps2histo.py*; for details on the procedure see https://github.com/wrf/lavaLampPlot). In contrast to the conceptually similar approach Blobtools [61] we used raw reads instead of contigs to yield a high resolution plot of read counts per %GC and 31mer coverage. The plot identified two read clouds with high counts at a kmer coverage of 80-140x (heterozygous cloud) and 160-260x (homozygous cloud), respectively. Additional read clouds at 270-320x and 380-41 Ox coverage mark repetitive sequence stretches. Another read cloud was found at a low coverage of 20-50x. Reads within this cloud and their pairs were extracted with *kmersorter.py* [added options: -s 0.16 -b 50 -w 0.40 -T -k 31] and *fastqdumps2histo.py*. Bowtie2 v2.2.5 [62] [added options: -q --no-sq] was used to map the 580,092 extracted reads to the 19 previously identified bacterial contigs (see section ‘Bacterial Contigs’ above). More than 86% of these reads mapped to the bacterial contigs confirming the bacterial origin of the reads within the low coverage read cloud. Read counts identified a relatively high abundance of bacterial cells and the GC content was similar to the host genome.

To estimate the per base genome coverage paired-end reads were mapped to the softmasked reference assembly with Bowtie2 v2.2.5 [added options: -q --no-unal --no-sq) and sorted with SAMtools vl.3.1 [63]. The bam file was used to create a bedgraph file in BEDtools v2.25.0 [64] by invoking the *genomecov* operation [added options: -ibam stdin -bga], A custom Python script (*bedgraph2histo.py*) [added options: -m 2000] was used to create a coverage histogram table. 81.4% of the genome falls within the second peak (165-332x coverage with a maximum at 248x) indicating that most of the genome was merged in the reference assembly (Figure S2).

### Genome completeness

#### Read and transcript mapping

To estimate the completeness of the reference assembly we first mapped paired-end reads and Moleculo reads back to the reference genome. For paired-end read mapping see section 7. above. BWA v0.7.12 [65] was used to map the Moleculo reads. Two successive rounds of mapping were performed with BWA *mem*. The first with stringent settings for long reads [added options: -k 200 -w 16000 -x intractg]. The output was filtered with the SAMtools v1.3.1 *view* script to receive mapped and unmapped reads. The 12,271 unmapped reads were mapped again using lower stringency settings to account for lower sequence identity in intergenic regions [added options: -w 16000 -x intractg]. More than 93% of the Moleculo reads and 84% of paired-end reads mapped back to reference contigs indicating a highly complete reference genome assembly and a low miss-assembly rate. For RNA-seq read mapping with Tophat2 see above.

Trinity transcripts and transdecoder predicted protein coding sequences were mapped to the hardmasked genome with GMAP v2015-07-23 [66] [added options :-f 3 -B 5 -n 1 --cross-species]. All DNA, RNA and transcripts mapping stats are summarized in Table S3.

#### BUSCO gene set

To further evaluate genome completeness we screened for a set of presumptive single copy proteins conserved in all animals, the BUSCO gene set. BUSCO v2.0 [67] was run separately on the *de novo* (BRAKER1) proteins, the StringTie transdecoder proteins, and the Trinity transdecoder proteins [added options: -1 metazoa_odb9 -m prot], respectively. It identified between 89-93% complete proteins (Table S4) indicating an almost complete reference genome and transcriptome.

### Synteny

To identify collinearity between the two placozoan species all *X. yyyyyyyyyyyyy* contigs >100kb were aligned to the largest ten *T. adhaerens* scaffolds (accounting for 70.3Mb or 66.5% of the genome assembly; including 5.7Mb gaps) with default settings. For the alignments the LASTZ vl.02.00 [68] (implemented as plugin in Geneious) was used. Of the 222*X. yyyyyyyyyyyyy* contigs >100kb a total of 144 (accounting for 60.6Mb or 69.4% of the genome assembly) aligned to the ten largest *T. adhaerens* scaffolds. Aligned *X. yyyyyyyyyyyyy* contigs were extracted from the assembly, sorted and occasionally reverse complemented to be oriented according to the *T. adhaerens* scaffolds. Gene annotations (GFF) of contigs as well as protein sequences were extracted for the target scaffolds/contigs sets of both species. A MCScanX run [69] [added option: -a] was performed for each target set using the extracted *T. adhaerens* and *X. yyyyyyyyyyyyy* GFF’s together with the reciprocal best five BLASTP hits [added options: -evalue 1e-10 -max_target_seqs 5 -outfmt 6] between and among proteins of both placozoans. Dual synteny line plots of the resulting collinearity files were visualized in VGSC v1.1 [70] [added options: -tp DualSynteny] and combined to Figure 4 in EazyDraw v3.10.3 (Dekorra Optics, LLC enterprises). In addition bar plots were generated for the ten *T. adhaerens* scaffolds and and the matching 144 *X. yyyyyyyyyyyyy* contigs in VGSC [added option: -tp Bar], Bar plots were mapped onto the DualSyntheny plots to show collinearity within each set and macrosynteny between both genomes. The percentage of collinearity between the *T. adhaerens* scaffolds and *X. yyyyyyyyyyyyy* contigs was calculated in MCScanX and results for the ten scaffolds are given in Table S4. The mean collinearity was calculated as the sum of the individual collinearities for the ten *T. adhaerens* scaffolds multiplied by a size correction faction for each scaffold (i.e. percent coverage of the totally evaluated 70.4Mb of the *T. adhaerens* genome).

Syntenic block sizes and number of blocks were calculated using the custom Python script m*icrosynteny.py* (described in [71]) with skipping no more than one gene [added option: -s 1] and otherwise default options.

### Intraspecific sequence variation

#### Genomic SNPs

Single nucleotide polymorphisms (SNPs) were identified with two independent tools, FreeBayes v0.9.21 [72] and GATK v3.5 [73,74]. For both analyses the bam file of Bowtie2 mapped reads (see section 7.) was used as input.

FreeBayes was run in parallel mode and the resulting vcf file was filtered with VCFfilter [added options: -f "QUAL > 20"]. For the GATK analysis the GATK best practice guidelines for variant discovery in DNAseq was followed (https://software.broadinstitute.org/gatk/best-practices/). Initially an index of the reference contigs was generated with SAMtools and a dictionary file with the Picard Tools v 2.3.0 *CreateSequenceDictionary* script (http://broadinstitute.github.io/picard). Read groups were then defined, reads sorted, duplicates marked and an index created with the Picard Tools scripts *AddOrReplaceReadGroups* [added options: SO=coordinate) and *MarkDupIicates* [added options: CREATE_INDEX=true, VALIDATION_STRINGENCY=SILENT, M], Processes files were used for the successive GATK variant calling using a set of scripts. Base frequencies were recalibrated with *BaseRecalibrator* [added options: -net 8, -knownSites] using the FreeBayes vcf as recalibration input. A second pass was run using the produced recalibration table to analyze covariation remaining after recalibration. *AnalyzeCovariates* [added options: -before, -after, -plots] was used to generate a comparison plot of reads before and after recalibration (not shown). As recalibration improved read quality scores the recalibration was applied to the sequence data with *PrintReads* [added options: -net 8, -I, -BQSR], Variants were then called using the recalibrated reads with *HaplotypeCaller* [added options: -net 8, --genotyping_mode DISCOVERY, -stand_call_conf 10 -stand_emit_conf 30], SNPs were extracted from the call set with *SelectVariants* [added options: -selectType SNP], Highly stringent SNP filtering was performed with *VariantFiltration* [added options: --filterExpression "QD < 2.0 ‖ FS > 60.0 ‖ MQ < 40.0 ‖ MQRankSum < −12.5 ‖ ReadPosRankSum < −8.0"]. Indels were extracted from the variant call set with *SelectVariants* [added options: -selectType INDEL] and filtered with *VariantFiltration* [added option: --filterExpression "QD < 2.0 ‖ FS > 200.0 ‖ ReadPosRankSum < -20.0"]. This procedure identified 1,397,488 high confidence genomic SNPs in the *X. yyyyyyyyyyyyy* DNA equaling roughly 16 SNPs per 1kb or a heterozygosity of 1.6%.

To identify SNP in the exonic, intronic and intergenic fraction of the genome the FreeBayes vcf (see section above) was input in a custom Python script (*vcfstats.py*) together with the stringtie annotation gtf and the stringtie transdecoder annotation GFF file (see section ‘StringTie gene models’ below for details). A plot of the SNP numbers against the coverage identified the heterozygous and homozygous peaks with differences in SNPs between the genomic fractions (Figure S3). The exonic fraction showed almost no SNPs within the heterozygous and the highest number in the homozygous peak, whereas the intergenic fraction had a larger number of SNPs in the heterozygous and a reduced number in the homozygous peak. The intronic fraction is an intermediate of the two. This indicates that (1.) most of the genic (exonic & intronic) regions have been successfully merged in the assembly process resulting in an almost completely merged reference assembly, and (2.) the proportion of unmerged haplocontigs is essentially higher in the intergenic fraction. This confirms an expected higher sequence divergence between the two genomic haplotypes in intergenic regions.

#### SNPs in RNAseq data

To call RNAseq variants the GATK best practice guidelines for variant calling on RNAseq was followed [74,75]. The Tophat2 RNAseq mapping bam file (see above) was used. The index and dictionary files were generated as for DNA SNPs (above). Read groups were defined, reads sorted, duplicates marked and an index created with the Picard Tools as mentioned. Process files were used for the successive GATK variant calling using a set of scripts. To split reads into exon segments, hard-clip any sequences overhanging into the intronic regions and to reassign mapping qualities the *SplitNCigarReads* script was applied [added options: -rf ReassignOneMappingQuality -RMQF 255 -RMQT 60 −U ALLOW_N_CIGAR_READS], Base recalibration (one round) and read printing were performed as for DNA. Variant calling of recalibrated reads was done with *HaplotypeCaller* [added options: -dontUseSoftClippedBases -stand_call_conf 10.0 -stand_emit_conf 30.0] and stringent filtering with *VariantFiltration* [added options: -filterName FS -filter "FS > 30.0" -filterName QD -filter "QD < 2.0"]. This procedure identified 302,430 high confidence SNPs in *Xxxxxxxxx yyyyyyyyyyyyy* strain M153E-2 RNAseq data.

#### Comparison of genomic and transcriptomic SNPs

SNP numbers and sites were compared between the two *Xxxxxxxxx* strains. First, all identified DNA and RNA SNPs within predicted BRAKER1 exons were extracted separately with BEDtools *intersect* [added options: -a -b -wa]. Second, unique DNA and RNA SNPs were extracted with BEDtools *intersect* [added options: -a -b -v -f 1.0 -wa]. This procedure identified a total of 138,302 (45.7% of all) RNA SNPs in exons 21,963 (15.7%) of which are unique to strain M153E-2. This is the equivalent of one unique SNP per kilobase coding sequence. In contrast a total of 202.901 (14.5% of all) DNA SNPs were identified in exons in the DNA strain with 86,278 (6.2%) unique exonic SNPs or 4 SNPs per kilobase coding sequence. Combined SNP counts indicates very low differences between the two strains with only 0.5% unique SNPs in coding sequence. The number for intronic regions is expected to be higher but as no genomic data is available from M153E-2 this cannot be tested. All SNPs counts are summarized in Table S2.

### Intra- and interspecific placozoan distances

#### Full-length allele identification

To identify all loci for which both full-length alleles were available we extracted reference gene sequences (CDS and introns) plus 1kb sequences upstream and downstream based on the BRAKER1 annotation GFF file. Only the longest gene model was used for each gene. Haplocontigs generated by SPAdes (first step in the dipSPAdes assembly pipeline) were mapped against the extracted reference gene sequences with BWA mem [added options: -k100 -W40 -rl0 -A1 -B1 -O1 -E1 -L0], Unmatched regions of the haplocontigs were hard clipped with Bamutils *removeclipping* of the NGSUtils v0.5.7 [76] with default settings. This also trimmed the overhanging haplocontigs to the reference sequence length. After a size filtering of mapped contigs with Bamutils filter [added options: -minlen 1000] the bam file was sorted with SAMtools view. All alignments were loaded into Geneious R8 and filtered to keep only loci with (1) 100% reference coverage, (2) exactly two mapped haplocontigs, and (3) both haplocontigs spanning the BRAKER1 gene model in the reference. This resulted in 5,401 loci for which the reference and both allele sequences were extracted and gaps removed.

#### Reference and allelic gene models

To identify CDS in alleles of all loci we performed RNA mapping to the three sequences for each of the 5,401 loci with Tophat2. The BRAKER1 pipeline was then run with the generated RNA-mapping bam file with changes in some BRAKER1 scripts: (1) ' --min_contig=100' was added to GeneMark-ET command line (1. 616) to perform training on contigs with at least lkb (instead of 50kb), and (2)'alternatives-from-evidence=$alternatives_from_evidence' replaced by '--genemodel=exactlyone' in the BRAKER1 script to predict only one gene for each allelic contig. Coding sequences from the BRAKER1 predictions were extracted and assembled in Geneious allowing for 20% sequence difference, 20 % gaps, 500bp gap size and multiple mapping. Loci with more or less than three sequences were excluded from further analyses. This resulted in 4,452 loci with full-length gene models in exactly three sequences (reference, allele A, and allele B).

#### Trichoplax adhaerens *orthologs identification*

To identify orthologs we built Hidden Markov Models with HMMER for the 4,452 loci using protein translations (created with the custom Python script *prottrans.py*) of the three *X. yyyyyyyyyyyyy* coding sequences each. We also used these proteins to create reference BLAST databases. HMMs and BLAST databases were used to identify orthologs using HaMStR v13.2.6 [77] [added options: -est-eval_BLAST=le-30 -eval_hmmer=le-30 -strict], A total of 3,984 orthologs were identified in *T. adhaerens* using the stringent HaMStR search.

#### Ortholog refinements

To further refine ortholog predictions and to remove false positives orthologs, reciprocal blast searches (evalue le-10) of the 3,984 HaMStR orthologs were performed for *X. yyyyyyyyyyyyy* and *T. adhaerens*. We furthermore performed blast searches (evalue 1e-10) of each ortholog set against the human uniprots. Based on the blast results we kept only those orthologs that (1.) resulted in reciprocal best hits between the two placozoans and (2.) had the same best blast hit to the human uniprot protein. This stringent procedure further reduced the final ortholog set to 2,720 very likely true orthologs.

#### Alignments and distance calculations

Protein sequences were aligned for each *X. yyyyyyyyyyyyy* and *T. adhaerens* for all 2,720 protein pairs with ClustalO vl.2.0 [78] [added options: --percent-id --full --output-order=input-order]. The nucleotide CDS were back-aligned based on the untrimmed protein alignment using a custom Python script (*regapper.py*). Interspecific (*X. yyyyyyyyyyyyy* allele A vs B) and intraspecific (*X. yyyyyyyyyyyyy* reference vs *T. adhaerens*) distances were calculated in ClustalO [added options: --percent-id --full --output-order=input-order --distmat-out=outfile] based on the Gblocks trimmed CDS and protein alignments.

### dN/dS ratios and codon saturation

Ratios of non-synonymous to synonymous substitutions (dN/dS) as well fractions of unchanged codons, synonymous and non-synonymous sites were calculated based on a custom Python script (alignmentdnds.py) using re-gapped CDS alignments and untrimmed protein alignments (Figure S4). Codons with any ambiguous bases and gapped sites were ignored.

### Genetic distances in non-bilaterians

To estimate molecular differences between *X. yyyyyyyyyyyyy* and *T. adhaerens* and to bring these into a taxonomic context we measured genetic distance using an extended data matrix of 212 nuclear proteins set up by Cannon *et al*., 2016. This data matrix was chosen as it includes a comparable number of sites for a diverse taxonomic range and is therefore also suitable for phylogenetic analyses. In addition genetic distances were measured for five standard barcoding (‘selected’) markers, namely nuclear ribosomal subunits 18S (Figure S6) and 28S (Figure S7), mitochondrial large ribosomal subunit 16S (Figure S8), as well as the mitochondrial proteins COl (Figure S9) and ND1 (Figure S10). An overview of means for all distances of all six marker sets is provided as Figure S11. The incorporation of datasets from four individual categories (nuclear protein vs. nuclear ribosomal DNA vs. mitochondrial protein vs. mitochondrial ribosomal DNA) enabled the comparison among markers with individual substitution rates.

#### Ortholog identification and alignment of nuclear proteins

Orthologs of the 212 genes were identified for *Xxxxxxxxx yyyyyyyyyyyyy*, *Trichoplax adhaerens* as well as a set of selected sponges, cnidarians and ctenophores (see Table S6 for a list of used taxa) using HaMStR. Transcriptomes were either downloaded from respective sources or, if no transcriptome was available, an assembly was generated with Trinity v2.0.6 [added options: --normalizereads --trimmomatic]. All used transcriptomes were translated using a custom Python script (*prottrans.py*) keeping only proteins with at least 50 amino acids [added options: -r -m -n -a 50], To perform ortholog searches Hidden Markov Models (HMMs) were built for all genes based on the final Cannon *et al.* protein alignments with HMMER. Using the sequences included in their alignments, reference BLAST datasets were created for the two outgroups (*Monosiga brevicollis*, *Salpingoeca rosetta*), all non-bilaterians (*Trichoplax adhaerens, Amphimedon queenslandica, Leucosolenia complicata, Aphrocallistes vastus, Oscarella carmela, Craspedacusta sowerby, Nematostella vectensis, Stomolophus meleagris, Euplokamis dunlapae, Mnemiopsis leidyi, Pleurobrachia bachei*), plus *Drosophila melanogaster* and *Homo sapiens*. The first HaMStR run was performed on the translated unigenes of a limited broad range taxon set, which included representatives from all non-bilaterian phyla and all classes within these, where available. In this first run all 15 reference taxa mentioned above were used [added options: -eval_hmmer=le-10 -eval_BLAST=le-10 -representative -append -strict], HaMStR outputs were transformed to fasta format and redundant orthologs of the 15 HaMStR runs for each proteome were filtered with a custom Python script (*commonseq.py*) [added options: -t p]. Sequences of individual ortholog groups (OG) for all taxa were combined to separate fasta files, which were aligned with the respective untrimmed alignment (kindly provided by Johanna Taylor Cannon) using MAFFT v7.273 [79] [added options: -linsi --amino --leavegappyregion]. Trimmed sequences from the Cannon *et al.* 212 gene set were aligned to the first alignment again with MAFFT and the same options. This procedure enabled accurate alignment of the trimmed sequences with the newly added sequences. The second alignment was trimmed according to the included trimmed sequences and used to create a second set of HMMs and BLAST reference taxa for another HaMStR run on the remaining proteomes. In this run we used *Monosiga brevicollis*, *Salpingoeca rosetta, Drosophila melanogaster*, *Homo sapiens* as core reference taxa plus an individually selected set of reference taxa for the four non-bilaterian phyla: (1.) each one taxon of the Anthozoa, Hydrozoa, Scyphozoa, and Cubozoa for Cnidaria, (2.) each one reference taxon of the Calcarea, Hextactinellida, and Homoscloromorpha as well as two of the Demospongiae for Porifera, (3.) *Pleurobrachia bachei* and *Mnemiopsis leiydi* for Ctenophora, and (4.) *Trichoplax adhaerens* for Placozoa. Added options in the second HaMStR run were identical to the first and final alignments for orthologs were generated as stated before.

We carefully curated every single protein by generating single gene trees to identify contaminations and paralogs in the original Cannon *et al.* 212 protein dataset as well as in the newly added data. Filtering of paralogs was performed in PhyloTreePruner [80] based on trees generated with FastTree v2.1.5 [81] using default settings.

Based on this approach we identified a high rate of host (fish) contamination in several parasitic as well as (prey and algae) contaminations in two free-living cnidarians. The following taxa (Genbank accessions in parentheses) were excluded and are therefore not listed in Table S6: *Myxobolus cerebralis* (SRP045736), *Myxobolus pendula* (SRP063943), *Kudoa iwatai* (SRP042325), *Thelohanellus kitauei* (SRP020474), *Polypodium hydriforme* (SRP042947), *Platygyra carnosus* (accession: SRP010342) and *Podocoryne carnea* (SRP041583).

After pruning, alignments were inspected manually and miss-aligned sequence ends were trimmed to the next unambiguously aligned position with respect to the next closest related taxa. Finally, Operational Taxonomic Units (OTUs) were generated for closely related taxa to increase the matrix densities for closely related taxa by merging the protein sets (see Table S7). The final alignment is referred to as dataset 1 (Table S8).

This two stage HaMStR approach using a broad phylogenetic range of reference taxa in the first and multiple selected taxa in the second run resulted in a higher yield of orthologs compared to a single run with a single and distantly related taxon (e.g. *Drosophila melanogaster*) alone.

#### Orthologs identification and alignment of selected barcoding markers

Mitochondrial markers were extracted from public mitochondrial genomes if available (Table S10). To retrieve mitochondrial genes from taxa without published mitochondrial genomes we performed BLASTN/TBLASTX (evalue IE-5) searches against available transcriptomes (Table S6). Nuclear ribosomal DNA (rDNA) sequences were identified by BLASTN searches against transcriptomes using the rDNA sequence of the next closest related taxa for which sequence information was available. For all included Porifera, Cnidaria and Ctenophora taxa we could isolate full-length 18S and 28S sequences from transcriptomic/genomic data and in most cases even the full-length rDNA cascade (including ITS1/2 and 5.8S). We used the placozoan rDNA accessions AY652583.1, AY652578.1, AY652585.1, AY652580.1, AY652587.1, AY652581.1.

Multiple sequence alignments were generated with MAFFT using the LINSI algorithm for protein sequences (CO1, ND1) and the GINSI algorithm for ribosomal genes (16S, 18S, 28S) with otherwise default settings. Individual alignments were created for each class within Porifera and Cnidaria to reduce unambiguously aligned sites. For the Placozoa and Ctenophora we used all sequences to generate a single alignment for each marker.

#### Distance calculations

Mean group pairwise genetic distances were calculated in MEGA7 [82] [settings: model/method=p-distance; gaps/missing=pairwise]. Groups were assigned to all taxa and between group mean distances were calculated for orders within classes, families within orders and genera within families for the non-bilaterian phyla Porifera, Cnidaria and Ctenophora. The nuclear protein distance in placozoans was calculated for *T. adhaerens* and *X. yyyyyyyyyyyyy* only, since no other genomes are available.

To calculate genetic distances of selected single markers within the Placozoa two additional undescribed placozoan species (sp. H4 and sp. H8) were included. These two taxa were included for a better representation of genetic distances within the entire phylum. According to the established placozoan 16S molecular phylogeny [6], *Xxxxxxxxx yyyyyyyyyyyyy* and Placozoa sp. H4 represent closely related taxa within placozoan subgroup A2, Placozoa sp. H8 represents subgroup A1 and *T. adhaerens* represents group B.

### Phylogenetic trees

To assess the effect of adding a second placozoan species on the placement of the Placozoa in the animal tree of life and to estimate branch length to the two placozoan species, dataset 1 was further condensed to generate a highly complete protein matrix (dataset 2) with only 10.8% missing characters in 58 taxa, including 32 non-bilaterians and two outgroups (both with sequence information for all 212 proteins).

It has been clearly demonstrated that the CAT model (specifically CAT-GTR) implemented in PhyloBayes [83] fits phylogenomic amino acid super-matrices containing non-bilaterians best [30,84]. However, the computational burden of reaching convergence of analyses using the CAT-GTR model can be prohibitive. It is also well known that phylogenomic datasets frequently suffer from compositional heterogeneity that might negatively influence phylogeny estimation [85–87]. Compositional heterogeneities can be reduced by the so-called “Dayhoff recoding” [34,88,89] which combines amino acids with similar physicochemical properties into one of six categories. Through this reduction of character space, lineage-specific compositional heterogeneities are lessened, at the cost, however, of losing phylogenetic signal (e.g. [35]). However, another advantage of Dayhoff recoding is a significant reduction of computation time needed to reach convergence.

The protein as well as the Dayhoff 6-state recoded dataset 2 were analysed with PhyloBayes MPI vl.7 [36,83], employing the CAT-GTR model, on the Linux cluster of the Leibniz Rechenzentrum (www.lrz.de) in Garching bei München, running two chains (each on 104 CPUs) each until reaching convergence, as estimated by using *tracecomp* and *bpcomp* programs of the PhyloBayes package (see PhyloBayes manual for details). Phylogenetic trees are shown as Figures 4, S12 and S13.

### Data and software availability

Raw genomic short and long reads as well as RNAseq reads, respectively, have been deposited at NCBI Short Read Archive under SRRxxxx, SRRxxxx, SRRxxxx. Bioproject accession is PRJNAxxxx.

A repository has been created that hosts all files related to the genome and performed analyses (http://bitbucket.org/molpalmuc/XXXXX):

- Masked and unmasked reference genome assembly [fasta]
- Transcriptome and proteome versions [fasta]
- Annotation tracks [GFF]: genes, CDS, mapped transcripts, SNPs, unexpressed ab initio gene models, tRNAs, repeats
- Raw coding sequences for 2,720 orthologs [fasta]
- Raw protein sequences for 2,720 orthologs [fasta]
- Raw coding sequences alignments for 2,720 orthologs [fasta]
- Raw protein sequences alignments for 2,720 orthologs [fasta]
- Trimmed coding sequences alignments for 2,720 orthologs [fasta]
- Trimmed protein sequences alignments for 2,720 orthologs [fasta]
- Protein matrix used for distance calculations (dataset 1) [phylip]
- Alignments of selected single marker sequences for distance calculation (16S, coxl, nadl, 18S, 28S) [fasta]
- Protein alignments for 212 proteins used for phylogenetic inferences [fasta]
- Protein matrix used for phylogenetic inferences (dataset 2) [phylip]
- Dayhoff 6-state recoded protein matrix used for phylogenetic inferences (recoded dataset 2) [phylip]
- Partition files for dataset 1 & 2
- Output files from phylobayes analyses of protein and Dayhoff 6-state recoded dataset 2

Python scripts used in this study are available at https://bitbucket.org/wrf/sequences and https://github.com/wrf/lavaLampPlot.

## Footnotes

**NOTE RELATING TO TAXONOMIC RULES:** According to the <u>International Code of Zoological Nomenclature</u> preprint publication of taxonomic names is discouraged. Consequently, the “*Xxxxxxxxx yyyyyyyyyyyyy* / *X. yyyyyyyyyyyyy*” given here is a dummy only. The valid name will be available upon formal journal publication.

